# Biomolecular Contrast Agents for Optical Coherence Tomography

**DOI:** 10.1101/595157

**Authors:** George J. Lu, Li-dek Chou, Dina Malounda, Amit K. Patel, Derek S. Welsbie, Daniel L. Chao, Tirunelveli Ramalingam, Mikhail G. Shapiro

**Author notes:** Correspondence should be addressed to M.G.S., T.R., or D.L.C.

## Abstract

Optical coherence tomography (OCT) has gained wide adoption in biological and medical imaging due to its exceptional tissue penetration, 3D imaging speed and rich contrast. However, OCT plays a relatively small role in molecular and cellular imaging due to the lack of suitable biomolecular contrast agents. In particular, while the green fluorescent protein has provided revolutionary capabilities to fluorescence microscopy by connecting it to cellular functions such as gene expression, no equivalent reporter gene is currently available for OCT. Here we introduce gas vesicles, a unique class of naturally evolved gas-filled protein nanostructures, as the first genetically encodable OCT contrast agents. The differential refractive index of their gas compartments relative to surrounding aqueous tissue and their nanoscale motion enables gas vesicles to be detected by static and dynamic OCT at picomolar concentrations. Furthermore, the OCT contrast of gas vesicles can be selectively erased *in situ* with ultrasound, allowing unambiguous assignment of their location. In addition, gas vesicle clustering modulates their temporal signal, enabling the design of dynamic biosensors. We demonstrate the use of gas vesicles as reporter genes in bacterial colonies and as purified contrast agents *in vivo* in the mouse retina. Our results expand the utility of OCT as a unique photonic modality to image a wider variety of cellular and molecular processes.

## INTRODUCTION

Optical coherence tomography (OCT) occupies a unique niche as an optical imaging modality that can visualize biological structures and processes with the spatial resolution of microns, the temporal resolution of milliseconds and a penetration depth of 1-3 mm^1, 2^. Since its invention, OCT has found utility in a wide range of scientific and clinical applications, ranging from the study of bacterial biofilms to medical specialties such as ophthalmology, gastroenterology, cardiology, dermatology and pulmonology^3-9^. Despite this widespread use, OCT currently has a limited role in cellular and molecular imaging due to the lack of imaging agents capable of connecting OCT contrast to biological processes such as gene expression. In particular, while fluorescence microscopy takes advantage of synthetic fluorophores and genetically encoded reporters such as the green fluorescent protein (GFP) to visualize a wide variety of cellular functions, few widely used synthetic contrast agents and no reporter genes are currently available for OCT.

OCT typically relies on the backscattering of near-infrared (NIR) light by endogenous scatterers such as cell nuclei and mitochondria to resolve tissue morphology. To allow OCT to label and visualize specific molecules and cells within biological tissues^10-12^, previous work on OCT contrast agents has focused on synthetic particles such as gold nanorods^13-17^. While these materials produce strong NIR scattering, these synthetic agents must overcome delivery barriers to enter tissues and cells, may accumulate in the body after use, and are challenging to couple to intracellular processes such as gene expression. If OCT contrast agents could instead be made of genetically encodable biomolecules, they could serve as biodegradable, cell-produced imaging reagents and as reporter genes connected to specific gene expression circuits in engineered cells. However, despite progress in the development of genetically encoded contrast agents for other modalities^18, 19^, none exists for OCT.

Here we introduce the first biomolecular, genetically encodable OCT contrast agents. They are based on gas vesicles (GVs), air-filled protein nanostructures evolved in certain photosynthetic microbes as flotation devices to maintain optimal access to light and nutrients^20, 21^. GVs, which self-assemble inside the microbes based on a genetic program, comprise a hollow gas-filled interior with dimensions on the order of 200 nm, enclosed by a 2 nm-thick protein shell that is permeable to gas but excludes liquid water. As genetically encoded nanostructures, GVs can be expressed heterologously^22^ and have their properties modified through genetic engineering^23^. Recently, the mechanical and magnetic properties of GVs were utilized to develop molecular contrast agents for ultrasound^24^ and magnetic resonance imaging^25, 26^. Here we hypothesized that the differential refractive index of GVs’ air-filled interior relative to surrounding tissue would allow them to serve as genetically encoded reporters for OCT, and that their unique material properties would enable multiplexed and acoustically erasable molecular imaging. In this study, we set out to test these fundamental hypotheses through *in vitro* and *in vivo* experiments.

## RESULTS

### GVs produce OCT contrast

The refractive index of air at atmospheric pressure is 1.0, which differs substantially from the 1.3–1.4 values of most aqueous biological tissues (**Fig. 1a**). We reasoned the air contents of GVs would cause these nanostructures to strongly scatter incident light, allowing GVs-based contrast agents or cells expressing GVs to be visible on OCT images. To test the hypothesis, we acquired OCT images of hydrogels containing three different GV types derived from *Anabaena flos-aquae* (Ana GVs), *Halobacterium salinarum* NRC-1 (Halo GVs) and *Bacillus megaterium* (Mega GVs) (**Fig. 1b**). All three GV types produced significant OCT image contrast at nanomolar concentrations (**Fig. 1c**). Halo GVs, which have the largest average volume (**Fig. 1d** and **Supplementary Table 1**), produced the strongest scattering, followed by Ana GVs and Mega GVs, with respectively decreasing size (**Fig. 1e**). However, the scattering of each sample did not scale linearly with the total amount of gas encapsulated by GV particles (**Fig. 1f**), indicating a dependence on particle size and shape^27^. These results establish the basic ability of GVs to act as contrast agents for OCT and indicate that the properties of GVs could be tuned to modulate their contrast level.

**Fig 1.**
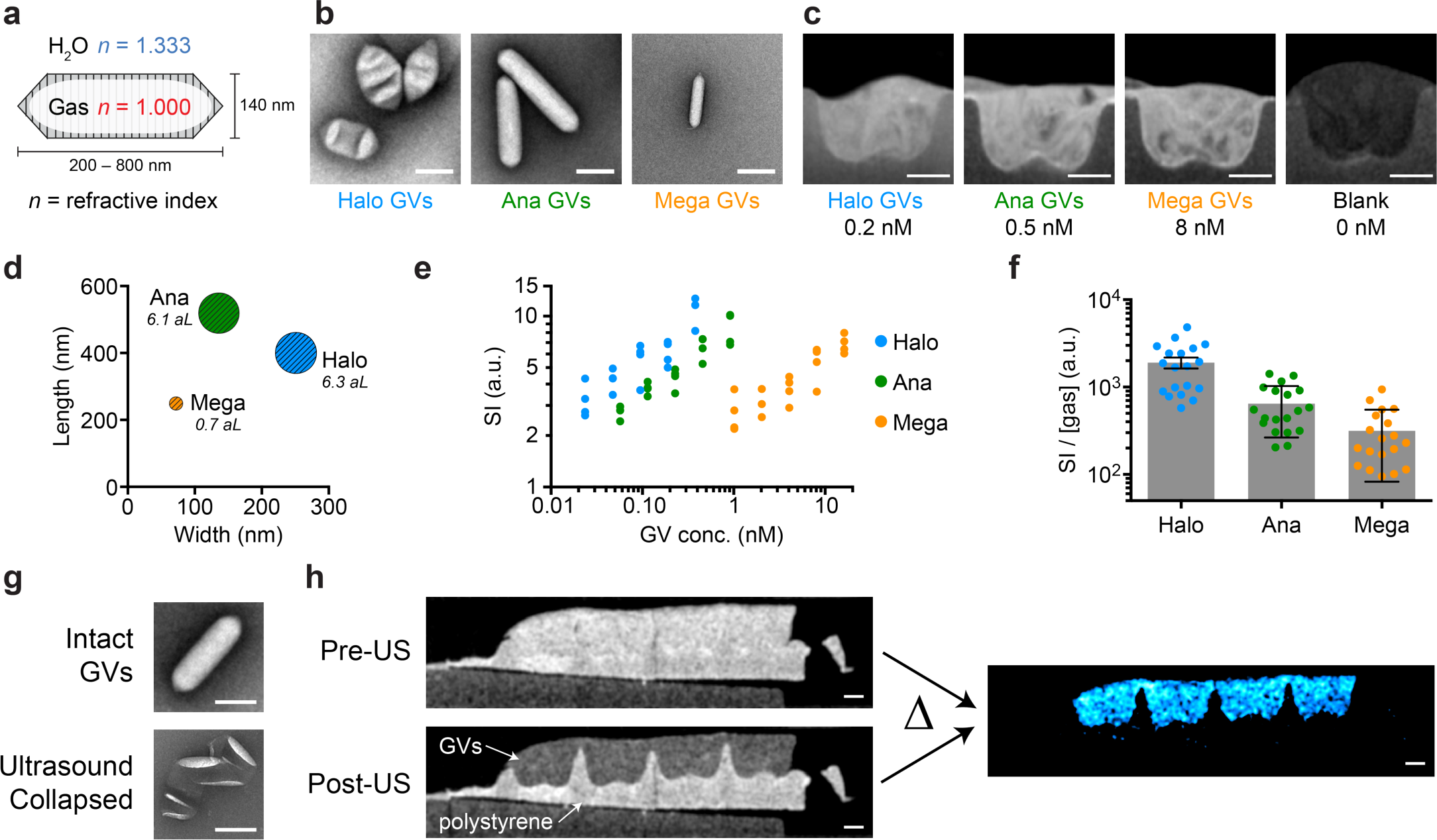
GVs produce OCT contrast that can be erased with ultrasound. **a**, Schematic drawing of a GV, the gas-filled interior of which has a refractive index (red) different from that of the surrounding H_2_O (blue). **b**, Representative transmission electron microscopy (TEM) and, **c**, OCT images of GVs from *Halobacterium salinarum NRC-1* (Halo), *Anabaena flos-aquae* (Ana) and *Bacillus megaterium* (Mega). GVs were embedded in a hydrogel phantom for OCT imaging. **d**, Mean length, width and gas volume (scaled to the areas of the symbols and labeled in attoliter) of Halo, Ana and Mega GVs. **e**, OCT signal intensity (SI) of the concentration series of Halo, Ana and Mega GVs. 𝒩 = 4 replicates. **f**, OCT SI normalized by the volume of gas contained inside GVs. Error bars represent SEM. **g**, Representative TEM images of Ana GVs before and after ultrasound treatment. **h**, OCT images of a phantom with regions containing either Ana GVs (OD_500,PS_ = 4) or polystyrene particles (mean diameter = 0.22 µm, 0.069% w/v, OD_500_ = 4), before and after the application of ultrasound. Subtracting these two images results in the difference image (colored in cyan). Scale bars represent 150 nm (**b**, **g**) and 0.5 mm (**c**, **h**). The dimensions of GVs in (**a**) are representative of Ana GVs.

### Acoustic erasure of GV contrast enables background subtraction

The detection of OCT contrast agents within tissues can be confounded by endogenous background scattering and speckle^12^. A common solution to mitigate this problem is to seek the “brightest” contrast agents that can surpass the contrast from a given tissue. However, this puts a stringent requirement on the design of contrast agents and often demands a relatively high concentration of agents delivered to the target tissue. In contrast, GVs have a unique built-in mechanism by which they can be distinguished from other scatterers. When GVs are subjected to pressure above a genetically determined threshold, they become irreversibly collapsed, leading to the rapid dissolution of their enclosed gas^20^, leaving behind just their protein shell (**Fig. 1g**). The required pressure can be delivered remotely and noninvasively using a brief pulse from an ultrasound transducer, allowing *in situ* GV collapse^25^. We therefore hypothesized that ultrasound pulses would erase GV-based OCT contrast, allowing it to be unambiguously distinguished from background. We tested this concept by imaging a hydrogel containing GVs and polystyrene nanoparticles as uncollapsible background scatterers (**Fig. 1h**). While it was impossible to identify the GV-containing region in the initial image, ultrasound treatment selectively erased the contrast from GVs, and subtraction of the pre- from the post-collapse image resulted in contrast specific to the GVs. Together with previous work showing that GV collapse does not affect cell viability^22^, these results suggest that acoustic collapse-based background subtraction is a powerful approach to enhancing OCT contrast specificity.

### Temporal characteristics of OCT images boost the detection sensitivity of GVs

Beyond static images, dynamic OCT provides information on the temporal characteristics of a sample. For example, during the time elapsed between two consecutive OCT images (usually several milliseconds), scatterers may re-orient relative to each other within, or move between, coherence volumes^10^. Brownian motion leads to dynamic changes in the speckle pattern commonly observed in OCT images of biological tissues, and has been used, for example, to measure anisotropic scatterer diffusivity^28^. Hypothesizing that the nanometer size of GVs would allow them to undergo significant Brownian motion, we wondered whether the temporal characteristics of OCT images would enable enhanced detection of GVs or contain information about their microscale arrangement in the sample.

To characterize the temporal characteristics of GV-based OCT contrast, we recorded a series of OCT images of GVs in hydrogel at 100 frames per second (fps) for 2 seconds. In GV samples that had an average signal intensity indistinguishable from a control polystyrene sample (**Fig. 2a**), the time trace of the signal intensity of individual pixels revealed significantly different temporal characteristics (**Fig. 2b**), which can be quantified as the pixel-wise temporal standard deviation (σ, **Fig. 2c**). Since the σ image provided a much stronger contrast than the average intensity (μ) image, we hypothesized that this image processing scheme can further boost the sensitivity of GV-based OCT contrast. Indeed, even our lowest tested concentration of Ana GVs (7.1 pM) showed a clear contrast with σ processing, which was not seen in a map of μ (**Fig. 2d** and **Supplementary Fig. 1**). These results establish the ability of GV-based contrast agents to be detected with boosted sensitivity using dynamic OCT.

**Fig 2.**
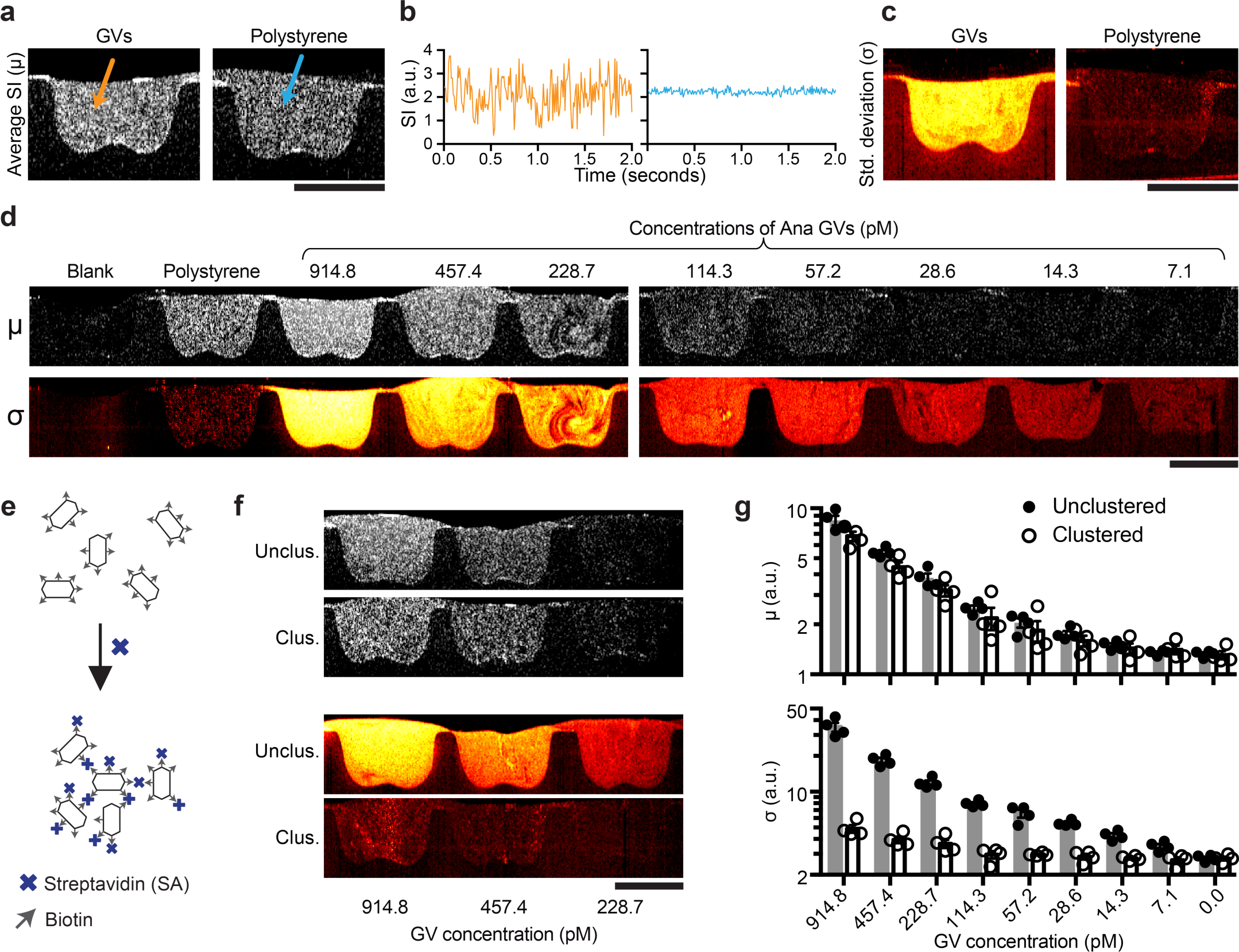
Temporal characteristics of OCT contrast of GVs and GV nanoassembly. **a**, Representative pixel-wise average signal intensity (SI) (µ image) of a series of OCT images collected over 2 seconds at 100 fps of Ana GVs (OD_500,PS_ = 4) and polystyrene particles (mean diameter = 0.22 µm, 0.069% w/v). **b**, Representative time traces of two individual pixels from region containing either GVs or polystyrene particles. **c**, Pixel-wise temporal standard deviation (σ image) of the same samples in (a). **d**, Representative µ and σ images of a concentration series of Ana GVs. **e**, Diagram of the clustering of surface-biotinylated Ana GVs by streptavidin. **f**, Representative µ and σ images of clustered and unclustered Ana GVs. **g**, Mean µ and σ. *N* = 4 replicates. Error bars represent SEM; scale bars represent 1 mm.

### Visualizing GV nanoassembly using dynamic OCT contrast

Having established the dynamic contrast behavior of independent GVs, we hypothesized that we could toggle their spatiotemporal dynamics and OCT contrast via clustering. This would enable the engineering of GV-based biosensors analogous to particle clustering sensors developed for other modalities^29-31^. We tested this hypothesis using surface-biotinylated Ana GVs mixed with tetrameric streptavidin as a model detectable analyte (**Fig. 2e**). The addition of streptavidin causes the GVs to form clusters of approximately 2 µm in hydrodynamic radius, ∼ 10 times larger than individual GVs. Upon clustering, the average OCT image intensity had a very minor change, while the temporal σσ decreased dramatically by about an order of magnitude (**Fig. 2, f-g** and **Supplementary Movie S1**). This shows that OCT contrast can track not only the concentration, but also the microscopic organization of GVs, allowing GVs to act as dynamic molecular sensors.

The decrease in dynamic contrast upon GV clustering is the expected result of a reduction in the number and movement of independent scatterers. Within the coherence volume, the number of individual GVs is on the order of 10^4^ to 10^6^ particles, within the optimal range for the phasor-summed multiple scattering susceptible to particle motion^10^. The clustering of GVs reduces the number of motionally independent scatterers to single digits, placing them outside the sweet spot for dynamic effects. In addition, clustering decreases the freedom of individual GV particles to undergo movements such as rotation. Since GVs are anisotropic in shape and may scatter light differently depending on their orientation^32^, restricting the reorientation of GVs is also expected to reduce the temporal fluctuation of their OCT contrast.

### OCT imaging of gene expression

Having established the OCT contrast of purified GVs, we next tested the ability of GVs to act as reporter genes in living cells. In particular, OCT has been used to visualize the structure bacterial biofilms^33^, an important class of living materials playing key roles in infectious disease, the human microbiome, environmental microbiology and synthetic biology^34-38^. Within this context, OCT offers a uniquely suitable combination of high spatial resolution, deep penetration and large field of view. To test the use of GVs as OCT reporter genes in bacteria, we used 3D volumetric OCT to image colonies of bacteria cultured on solid agar media. *E*. *coli* were engineered to express an 11-gene operon, ARG1, comprising a mixture of genes from Ana and Mega^22^. The expression of GVs was controlled using a *lac* operator and the chemical inducer isopropyl ß-D-1-thiogalactopyranoside (IPTG, **Fig. 3a**).

**Fig 3.**
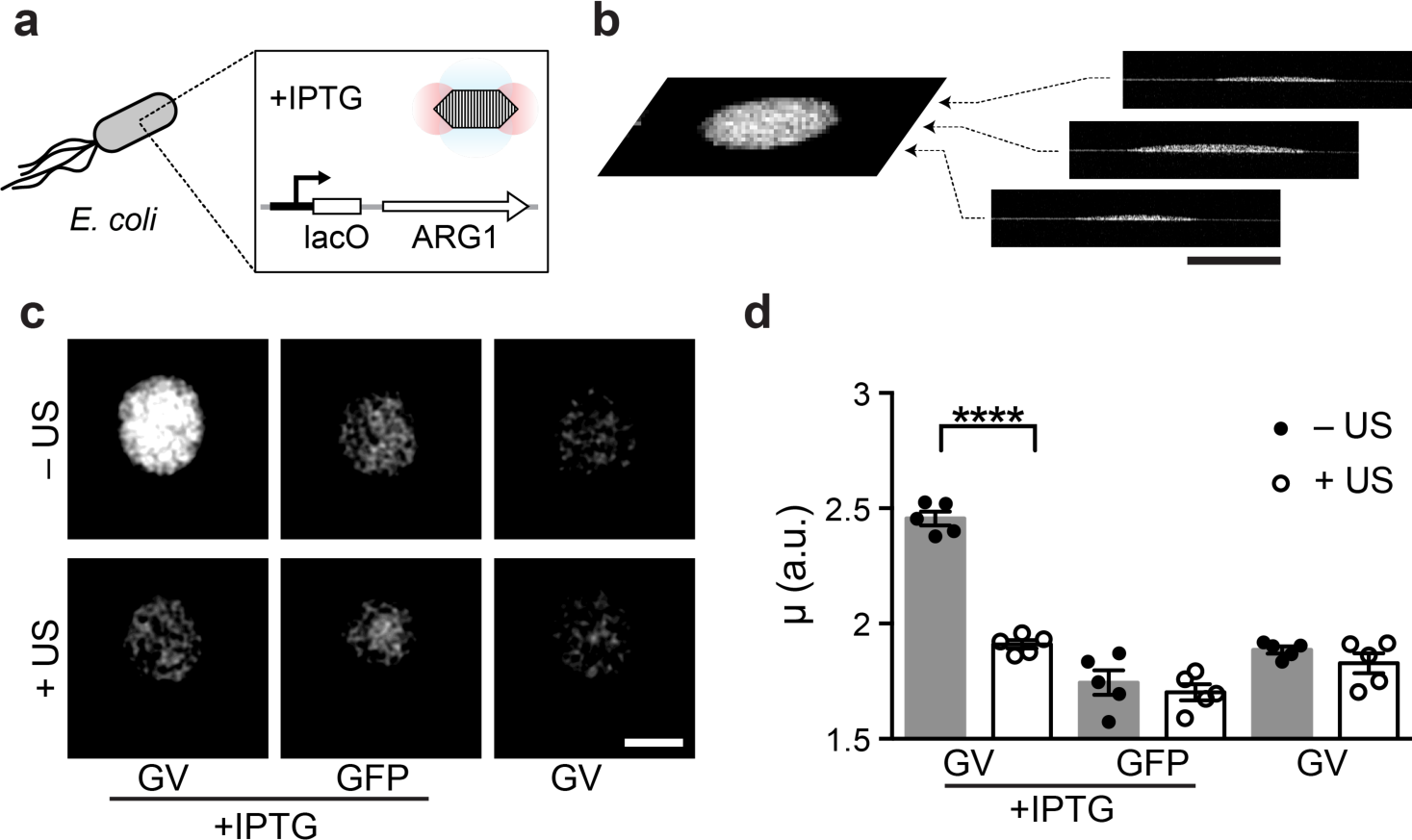
OCT reporter gene enables the imaging of gene expression in bacterial colonies. **a**, Diagram of the IPTG-inducible expression of ARG1 GVs inside *E*. *coli*. **b**, Representative coronal slice (C-scan image) extracted from 3D volumetric OCT image of an *E*. *coli* colony. **c**, Representative C-scans of colonies expressing GVs or green fluorescent proteins (GFP), in the presence or absence of the inducer, and subjected to ultrasound or left intact. **d**, Mean average intensity (µ) of *E*. *coli* colonies. 𝒩 = 5 biological replicates. Student’s t-test was performed between GVs with and without ultrasound treatment: *p* < 0.0001, unpaired, d.f. = 15.56. Error bars represent SEM; scale bars represent 1 mm.

Overnight growth on an agar plate containing IPTG allowed the formation of colonies expressing either ARG1 or GFP as a negative control for OCT contrast. The growth of ARG1 bacteria on a plate without IPTG served as additional, uninduced control. The shape of each colony was visible by 3D volumetric OCT (**Fig. 3b**), and GV-expressing colonies could clearly be distinguished from controls based on their enhanced contrast (**Fig. 3c**). After the ARG1 colonies were subjected to ultrasound pre-treatment to collapse the GVs, their contrast became indistinguishable from controls (**Fig. 3c**). Quantification of the OCT image intensity revealed that the expression of GVs made bacteria on average 36% brighter than controls (**Fig. 3d**). This indicates that GVs can function as OCT reporter genes in living cells.

### In vivo visualization of GVs in the retina

Finally, we tested the ability of GVs to be visualized by OCT within the context of living tissue *in vivo*. In particular, given the wide utility of OCT in ophthalmology^3^, and the need for molecular contrast agents to improve the diagnostic capabilities of this technique, we assessed whether GVs could be imaged in the eye. We injected GVs either intravitreally or into the subretinal space of the eye in anesthetized mice, and imaged the eye using an FDA-approved spectral domain OCT imaging device used in clinical ophthalmology (**Fig. 4a**).

**Fig 4.**
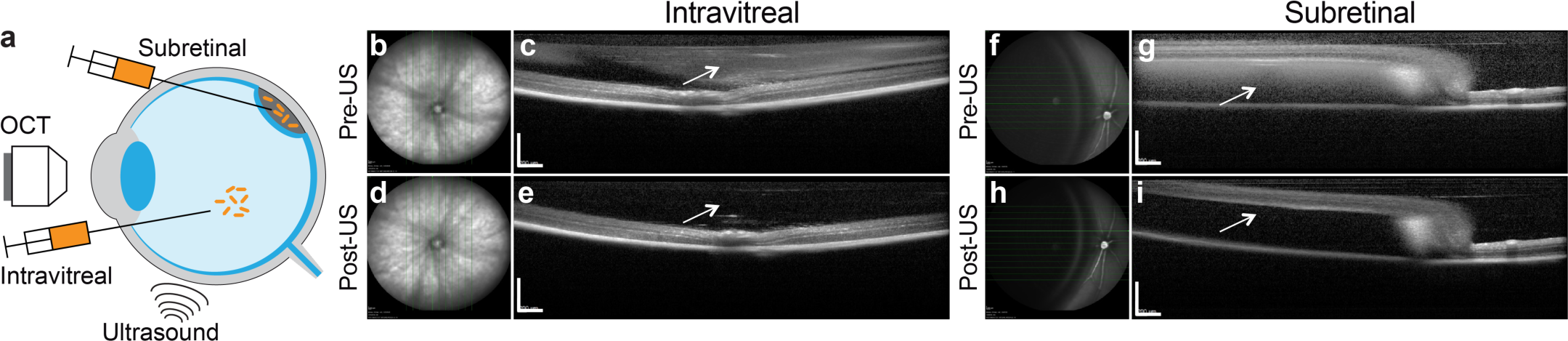
*In vivo* imaging of gas vesicles at mouse retina. **a**, Diagram of the experimental design. Objects are not drawn to scale. Representative infrared fundus image (**b**, **d**, **g**, **h**) and B-scan OCT images (**c**, **e**, **g**, **i**) of mouse eyes intravitreally (**b**, **c**, **d**, **e)** or subretinally (**f**, **g**, **h**, **i**) injected with 2 µl of 0.97 nM Ana GVs. Images of the same location of the eyes collected before (**b**, **c**, **f**, **g**) or after (**d**, **e**, **h**, **i**) ultrasound treatment. The arrows indicate the vitreous or subretinal space, where contrast from gas vesicles was expected. Scale bars represent 0.5 mm

GVs provided robust contrast in both the intravitreal (**Fig. 4, b-e**) and subretinal (**Fig. 4, f-i**) space. Furthermore, to unambiguously assign the observed contrast to injected GVs and test the feasibility of acoustic collapse *in vivo*, we delivered ultrasound pulses using a hand-held clinical transducer directly to the mouse eyes. Using the fundus images as a guide, we imaged the retina in approximately the same location to see the differences in OCT contrast before and after the ultrasound treatment, and found that the GV contrast was erased completely in both injected locations. These results were consistent across animal subjects (**Supplementary Fig. 2**, **a-b**). Likewise, if we injected GVs that were pre-collapsed by ultrasound *ex vivo*, there was no discernable change in OCT contrast (**Supplementary Fig. 2**, **c-d**). These experiments provide a proof of concept for the ability of GVs to serve as OCT reporters inside living animals.

## DISCUSSION

Our results establish GVs as the first biomolecular, genetically encodable OCT contrast agents. While this basic capability arises from the refractive index difference between GVs’ air interior and surrounding water, the contrast abilities of GVs are greatly enhanced by the fact that GVs are acoustically erasable and that their Brownian motion produces dynamic contrast fluctuation. This enables the unambiguous detection of GVs among background contrast and allows the development of GVs as functional sensors by coupling GV clustering to the presence of specific molecules.

The availability of biomolecular and genetically encodable OCT contrast agents may extend the use of OCT in new directions. For example, purified GVs have the potential to be developed as targeted molecular OCT contrast agents. GVs are biodegradable and readily amenable to genetic and chemical modification, allowing the addition of targeting moieties^23^. Targeted OCT contrast agents may find utility in combination with ophthalmic and intravascular OCT^39^. Furthermore, the genes encoding GVs could be incorporated into specific bacterial species, allowing their imaging within microbial communities, or the visualization of the engineered probiotics being developed as *in vivo* diagnostic and therapeutic agents. By connecting GV expression to conditional promoters, OCT contrast could further be used to visualize the dynamics of cellular function. Recent work to extend GV expression into mammalian cells^40^ could similarly enable the imaging of genetic and cellular therapies, such as those in development for inherited retinal degeneration^41^.

Building on these results, future studies can be designed to address current limitations and establish the wider applicability of GVs as genetically encodable OCT contrast agents. First, while the detection of GVs on OCT imaging of mouse retina demonstrates the feasibility of acoustically erasable OCT contrast *in vivo*, additional work needs to be done to realize the full potential of this technology, including the engineering of GVs for efficient penetration into tissues such as the retina^42, 43^ and an understanding of the biodegradation mechanisms of GVs. Secondly, the OCT contrast of GVs could be optimized through genetic engineering, since the size, shape and mechanical properties of GVs are fully genetically encoded. While the size and shape directly influence GVs’ scattering of light and Brownian motion, the mechanical properties of GVs will determine their acoustic collapse pressure threshold, providing a potential avenue towards multiplexed detection. Thirdly, the temporal characteristics of GV-derived OCT contrast require a more thorough and quantitative understanding^44^, which may be addressed through additional simulations and experiments. To this end, existing methods optimized to detect dynamic events such as polarization-sensitive OCT^45^ and Doppler OCT^46, 47^ may provide valuable information and better sensitivity in detecting GVs. These follow-up studies will increase the versatility and practicality of using GVs as biomolecular and genetically encodable OCT contrast agents, filling an important gap in this photonic technology’s unique ability to bridge spatiotemporal resolution and depth in biomedical imaging.

## MATERIALS AND METHODS

### Preparation, quantification and TEM imaging of GVs

The protocols to express and purify GVs were described previously^48^. Briefly, among the three genotypes of GVs used in this study, Ana GVs and Halo GVs were obtained from their native hosts, *Anabaena flos-aquae* (CCAP strain 1403/13 F) and *Halobacteria NRC-1* (Carolina Biological Supply), respectively, and Mega GVs were obtained by heterologously expressing the GV genes from *Bacillus megaterium* in *E*. *coli*. Following the harvest and lysis of the cells, GVs were purified through multiple rounds of centrifugally assisted floatation. Mega GVs, which are natively clustered after purification from bacteria, underwent additional steps of unclustering by 6M urea treatment, centrifugally assisted floatation and dialysis. The concentration of GVs was quantified by the pressure-sensitive optical density at 500-nm wavelength (OD_500,PS_). OD_500,PS_ was then converted to the molar concentration using the relation of 114, 47.3 and 2030 pM/OD_500,PS_ for Ana, Halo and Mega GVs, respectively (**Supplementary Table 1**). For TEM images, GVs were diluted to OD_500,PS_ ∼ 0.2 and then spotted on Formvar/Carbon 200 mesh grids (Ted Pella) that were rendered hydrophilic by glow discharging (Emitek K100X). After negative staining using 2% uranyl acetate, the samples were imaged on a Tecnai T12 LaB6 120 kV TEM.

### Clustering of GVs

Purified Ana GVs were biotinylated using EZ-Link Sulfo-NHS-LC-biotin (Thermo Fisher Scientific) according to the protocol described previously^48^, and the subsequent clustering was adapted from the previous procedure^25, 48^. Specifically, streptavidin (Geno Technology Inc.) was made into a stock solution of 5 mg/mL. 1 µL of the stock solution was added into every 100 µL of biotinylated Ana GVs at OD_500,PS_ = 16 and mixed immediately.

### Phantom preparation

Using a custom-3D-printed caster, a 1% agarose phantom was cast in phosphate-buffered saline (PBS) with hemispheric wells of diameter ∼ 2 mm. Room-temperature PBS containing various concentrations of GVs or polystyrene particles (0.069% w/v, OD_500_ = 4, mean diameter = 0.22 µm, Spherotech Inc.) were mixed quickly at 1:1 v/v with 2% agarose stock PBS solution maintained at 60 °C. 2 µL of the mixture was immediately loaded into a well, and the agarose solidified within a minute. Care was taken to avoid bubbles. To prepare for OCT imaging, the phantom was submerged in PBS solution.

### Bacterial colonies and ultrasound treatment

The plasmid pET28a_T7-ARG1 (Addgene plasmid no. 106473) was transformed into BL21(A1) one-shot competent cells (Thermo Fisher Scientific) for the expression of GVs, and GV genes in this plasmid were replaced by the mWasabi gene for the expression of GFP. Two-layer LB-Agar plates, each with 4 mm in thickness, were made according to a protocol adapted from previous literature^22^. The underlayer contained 50μg/ml kanamycin, 1.0% L-arabinose, and 0.8 mM IPTG. The overlayer contained 50 μg/ml kanamycin and 0.4% glucose. The overlayer was poured freshly and allowed to equilibrate to room temperature before the transformed cells were plated. Plates were then incubated at 30 °C for 24 h. To collapse the intracellular GVs in *E*. *coli*, ultrasound was applied through the bottom of the plates using a Verasonics Vantage programmable ultrasound scanning system equipped with the L11-4v 128-element linear array transducer using the following parameters: transmit frequency 6.25 MHz, transmit voltage 25 V, Tx focus = 10 mm and F = 0.1.

### OCT imaging and ultrasound treatment of purified gas vesicles and bacteria

The benchtop OCT scanning system used in this study was custom-built by OCT Medical Imaging Inc (OCTMI), and included a 1320 nm center wavelength 100 kHz swept laser (HSL-20-100M, Santec Corporation, Japan), a fiber optic interferometer, and a benchtop 2-axis galvo scanner setup. The detailed design, components and calculated resolution are described (**Supplementary Fig. 3**). For all the OCT images, A-lines were taken at 100 kHz rate with 512 points and an axial field of view of 4 ∼ 5 mm, and 1000 A-lines were included in each B-scan. For the concentration series of three types of GVs (**Fig. 1c-f**), each B-scan covered an image width of 14.6 mm, and in total 800 B-scans were collected while moving the imaging plane over a depth of 275 µm and averaged. For imaging the temporal characteristics (**Fig. 2**), the same image width was used, and 200 B-scans were collected without changing the position of the imaging plane. For bacterial colonies (**Fig. 3**), each B-scan covered an image width of 3.45 mm, and 1000 B-scans were collected while moving over a depth of 3.45 mm, thus allowing a square C-scan for each bacterial colony. To collapse GVs *in situ* (**Fig. 1h**), the portable Lumify Ultrasound System (Philips) was used with the following default settings: preset = MSK, depth = 3.5 cm, transducer = L12-4, power = 0.0 dB, MI = 1.0. The transducer was encapsulated with parafilm, submerged in water, and placed close to the sample perpendicular to the direction of OCT beams.

### Data processing

The raw OCT interferometric signal generated by the balanced detector was first digitized by a 12-bit, 500MS/sec waveform digitizer board (ATS9350, Alazar Technologies Inc., Canada) into a continuous data stream, which was subsequently stored in memory and transferred to the graphics processing unit (GPU) for real-time processing and frame-by-frame display. Each A-scan data array was window filtered, dispersion compensated, Fourier transformed, and the magnitude of the transformed data was calculated to generate depth-resolved OCT intensity data. Due to the exponential attenuation of OCT intensity data with respect to depth, logarithmic compression, thresholding and scaling were applied before the data could be mapped to an 8-bit bitmap image file for visualization and storage.

Fiji was used for subsequent processing and quantification of the image intensity^49^. These bitmap images were first converted from logarithmic scale back to linear scale, before applying any algorithms such as averaging over multiple frames (µ), image subtraction (Δ) or calculating standard deviation (σ) over a temporal image series. Afterwards, the regions of interest (ROIs) were selected, and the value of the average intensity in each ROI was obtained. Such values obtained in this linear scale were referred to as SI, µ or σ (**Fig. 1e**, **1f, 2b, 2g, 3d**). Afterwards, logarithmic compression was re-applied before the images were stored and displayed in the figures.

### In vivo OCT imaging and ultrasound treatment

All animal experiments were conducted in accordance with the Association for Research in Vision and Ophthalmology’s statement on the use of animals in ophthalmic research and were approved by the University of California San Diego Institutional Animal Care and Use Committee. For intravitreal and subretinal injections, 4-month-old C57B6/J mice (Jackson Laboratories) were anesthetized using a ketamine/xylazine cocktail injected intraperitoneally and eyes were locally anesthetized using 0.5% proparacaine hydrochloride eyedrop (Akorn, Lake Forest, IL). Mice remained anesthetized throughout the experiment. GVs were delivered either into the vitreous space through intravitreal injection or between the retina and retinal pigmented epithelium (RPE) through subretinal injection. 2 µl of 0.97 nM intact or collapsed Ana GVs in PBS (OD_500,PS_ = 8.5) were injected using 1.5 cm 33-gauge Hamilton needles (Hamilton Company, Reno, NV). A custom contact lens (plano power, black optic zone radius = 1.70mm, diameter = 3.2mm) (Cantor & Nissel Ltd, Brackley, UK) was placed on each mouse eye, and the eyes were dilated using 2.5% phenylephrine and 1% tropicamide eye drops (Akorn, Lake Forest, IL). OCT images were acquired using 11 B-scans in a 55 degree x 25 degree pattern (12.6 mm x 5.7 mm) by an HRA2/Heidelberg Spectralis (Franklin, MA). To collapse GVs inside mouse eyes, Lumify Ultrasound System (Philips) and the same parameters were used as described above for this device.

### Statistical Analysis

Sample sizes were chosen on the basis of preliminary experiments to give sufficient power for the reported statistical comparisons. Unless stated otherwise, statistical comparisons used two-tailed heteroscedastic t-tests with Welch’s correction.

## ACKNOWLEDGEMENTS

The authors thank Changhuei Yang for helpful discussion and Theodore Chang for assistance with initial experiments. Electron microscopy was performed at the Beckman Institute Resource Centre for Transmission Electron Microscopy at Caltech. This research was supported by the National Institutes of Health (grant numbers K99EB024600 to G.J.L.; K12EY024225 and P30EY022589 to D.L.C.; R44HL129496, R43HD071701 and R44CA177064 to T. R.; R01EB018975 to M.G.S.), the Defense Advanced Research Projects Agency (HR0011-17-2-0037 to M.G.S.), the Heritage Medical Research Institute (M.G.S.), the Packard Fellowship for Science and Engineering (M.G.S.), the Pew Scholarship in Biomedical Science (M.G.S.) and the Burroughs Welcome Fund Career Award at the Scientific Interface (M.G.S.).

## AUTHOR CONTRIBUTIONS

G.J.L., M.G.S., T.R. and D.L.C. conceived the research. G.J.L., M.G.S., T.R., L.C. and D.M. planned and performed the *in vitro* experiments, and G.J.L. and L.C. analyzed the data. D.L.C, A.K.P. and D.S.W. planned and performed the *in vivo* experiments using materials prepared by G.J.L. and D.M.. G.J.L. and M.G.S. wrote the manuscript with input from all authors.

## DATA AND MATERIALS AVAILABILITY

Raw data, gas vesicles and genetic constructs are available upon request to the authors.

## REFERENCES

1. Huang, D. et al. Optical coherence tomography. Science (New York, N.Y.) 254, 1178–1181 (1991).

2. Fujimoto, J.G. Optical coherence tomography for ultrahigh resolution in vivo imaging. Nat. Biotechnol. 21, 1361 (2003).

3. Hee, M.R. et al. Optical Coherence Tomography of the Human Retina. Archives of Ophthalmology 113, 325–332 (1995).

4. Tearney, G.J. et al. In vivo endoscopic optical biopsy with optical coherence tomography. Science (New York, N.Y.) 276, 2037–2039 (1997).

5. Izatt, J.A., Kulkarni, M.D., Hsing-Wen, W., Kobayashi, K. & Sivak, M.V. Optical coherence tomography and microscopy in gastrointestinal tissues. IEEE Journal of Selected Topics in Quantum Electronics 2, 1017–1028 (1996).

6. Liang, X. & Boppart, S.A. Biomechanical Properties of In Vivo Human Skin From Dynamic Optical Coherence Elastography. IEEE Trans. Biomed. Eng. 57, 953–959 (2010).

7. Fercher, A.F., Drexler, W., Hitzenberger, C.K. & Lasser, T. Optical coherence tomography-principles and applications. Rep. Prog. Phys. 66, 239 (2003).

8. Robles, F.E., Wilson, C., Grant, G. & Wax, A. Molecular imaging true-colour spectroscopic optical coherence tomography. Nature Photonics 5, 744 (2011).

9. Pahlevaninezhad, H. et al. Nano-optic endoscope for high-resolution optical coherence tomography in vivo. Nature Photonics 12, 540–547 (2018).

10. Yang, C. Molecular Contrast Optical Coherence Tomography: A Review. Photochem. Photobiol. 81, 215–237 (2005).

11. Boppart, S.A., Oldenburg, A.L., Xu, C. & Marks, D.L. Optical probes and techniques for molecular contrast enhancement in coherence imaging. JBO 10, 041208-041208-041214 (2005).

12. Boustany, N.N., Boppart, S.A. & Backman, V. Microscopic imaging and spectroscopy with scattered light. Annu. Rev. Biomed. Eng. 12, 285–314 (2010).

13. Liba, O., SoRelle, E.D., Sen, D. & de la Zerda, A. Contrast-enhanced optical coherence tomography with picomolar sensitivity for functional in vivo imaging. Sci Rep 6, 23337 (2016).

14. Oldenburg, A.L., Toublan, F.J.-J., Suslick, K.S., Wei, A. & Boppart, S.A. Magnetomotive contrast for in vivo optical coherence tomography. Opt. Express 13, 6597–6614 (2005).

15. Rao, K.D., Choma, M.A., Yazdanfar, S., Rollins, A.M. & Izatt, J.A. Molecular contrast in optical coherence tomography by use of a pump–probe technique. Opt. Lett. 28, 340–342 (2003).

16. Lee, T.M. et al. Engineered microsphere contrast agents for optical coherence tomography. Opt. Lett. 28, 1546–1548 (2003).

17. Mattison, S.P., Kim, W., Park, J. & Applegate, B.E. Molecular imaging in optical coherence tomography. Current molecular imaging 3, 88–105 (2014).

18. Lu, G.J., Farhadi, A., Mukherjee, A. & Shapiro, M.G. Proteins, air and water: reporter genes for ultrasound and magnetic resonance imaging. Curr. Opin. Chem. Biol. 45, 57–63 (2018).

19. Piraner, D.I. et al. Going Deeper: Biomolecular Tools for Acoustic and Magnetic Imaging and Control of Cellular Function. Biochemistry 56, 5202–5209 (2017).

20. Walsby, A.E. Gas vesicles. Microbiological Reviews 58, 94–144 (1994).

21. Pfeifer, F. Distribution, formation and regulation of gas vesicles. Nat Rev Micro 10, 705–715 (2012).

22. Bourdeau, R.W. et al. Acoustic reporter genes for non-invasive imaging of microorganisms in mammalian hosts. Nature 553, 86–90 (2018).

23. Lakshmanan, A. et al. Molecular Engineering of Acoustic Protein Nanostructures. ACS Nano 10, 7314–7322 (2016).

24. Shapiro, M.G. et al. Biogenic gas nanostructures as ultrasonic molecular reporters. Nat Nano 9, 311–316 (2014).

25. Lu, G.J. et al. Acoustically modulated magnetic resonance imaging of gas-filled protein nanostructures. Nature Materials 17, 456–463 (2018).

26. Shapiro, M.G. et al. Genetically encoded reporters for hyperpolarized xenon magnetic resonance imaging. Nat Chem 6, 629–634 (2014).

27. Jain, P.K., Lee, K.S., El-Sayed, I.H. & El-Sayed, M.A. Calculated Absorption and Scattering Properties of Gold Nanoparticles of Different Size, Shape, and Composition: Applications in Biological Imaging and Biomedicine. J. Phys. Chem. B 110, 7238–7248 (2006).

28. Marks, D.L., Blackmon, R.L. & Oldenburg, A.L. Diffusion tensor optical coherence tomography. Phys Med Biol 63, 025007 (2018).

29. Elghanian, R., Storhoff, J.J., Mucic, R.C., Letsinger, R.L. & Mirkin, C.A. Selective colorimetric detection of polynucleotides based on the distance-dependent optical properties of gold nanoparticles. Science (New York, N.Y.) 277, 1078–1081 (1997).

30. Perez, J.M., Josephson, L., O’Loughlin, T., Hogemann, D. & Weissleder, R. Magnetic relaxation switches capable of sensing molecular interactions. Nat Biotech 20, 816–820 (2002).

31. Lanza, G.M. et al. A novel site-targeted ultrasonic contrast agent with broad biomedical application. Circulation 94, 3334–3340 (1996).

32. Chhetri, R.K., Kozek, K.A., Johnston-Peck, A.C., Tracy, J.B. & Oldenburg, A.L. Imaging three-dimensional rotational diffusion of plasmon resonant gold nanorods using polarization-sensitive optical coherence tomography. PhRvE 83, 040903 (2011).

33. Wagner, M. & Horn, H. Optical coherence tomography in biofilm research: A comprehensive review. Biotechnol. Bioeng. 114, 1386–1402 (2017).

34. The Human Microbiome Project, C. et al. Structure, function and diversity of the healthy human microbiome. Nature 486, 207 (2012).

35. Flemming, H.-C. & Wingender, J. The biofilm matrix. Nature Reviews Microbiology 8, 623 (2010).

36. Liu, J. et al. Metabolic co-dependence gives rise to collective oscillations within biofilms. Nature 523, 550–554 (2015).

37. Danhorn, T. & Fuqua, C. Biofilm formation by plant-associated bacteria. Annu. Rev. Microbiol. 61, 401–422 (2007).

38. Lu, T.K. & Collins, J.J. Dispersing biofilms with engineered enzymatic bacteriophage. Proc. Natl. Acad. Sci. 104, 11197–11202 (2007).

39. Gora, M.J., Suter, M.J., Tearney, G.J. & Li, X. Endoscopic optical coherence tomography: technologies and clinical applications [Invited]. Biomedical Optics Express 8, 2405–2444 (2017).

40. Farhadi, A., Ho, G.H., Sawyer, D.P., Bourdeau, R.W. & Shapiro, M.G. Ultrasound Imaging of Gene Expression in Mammalian Cells. bioRxiv, 580647 (2019).

41. Trapani, I., Banfi, S., Simonelli, F., Surace, E.M. & Auricchio, A. Gene Therapy of Inherited Retinal Degenerations: Prospects and Challenges. Hum. Gene Ther. 26, 193–200 (2015).

42. Dalkara, D. et al. In Vivo–Directed Evolution of a New Adeno-Associated Virus for Therapeutic Outer Retinal Gene Delivery from the Vitreous. Science Translational Medicine 5, 189ra176–189ra176 (2013).

43. Lai, S.K., Wang, Y.-Y. & Hanes, J. Mucus-penetrating nanoparticles for drug and gene delivery to mucosal tissues. Adv. Drug Del. Rev. 61, 158–171 (2009).

44. Almasian, M., van Leeuwen, T.G. & Faber, D.J. OCT Amplitude and Speckle Statistics of Discrete Random Media. Scientific Reports 7, 14873 (2017).

45. Hee, M.R., Huang, D., Swanson, E.A. & Fujimoto, J.G. Polarization-sensitive low-coherence reflectometer for birefringence characterization and ranging. J. Opt. Soc. Am. B 9, 903–908 (1992).

46. Chen, Z., Milner, T.E., Dave, D. & Nelson, J.S. Optical Doppler tomographic imaging of fluid flow velocity in highly scattering media. Opt. Lett. 22, 64–66 (1997).

47. Izatt, J.A., Kulkarni, M.D., Yazdanfar, S., Barton, J.K. & Welch, A.J. In vivo bidirectional color Doppler flow imaging of picoliter blood volumes using optical coherence tomography. Opt. Lett. 22, 1439–1441 (1997).

48. Lakshmanan, A. et al. Preparation of biogenic gas vesicle nanostructures for use as contrast agents for ultrasound and MRI. Nature Protocols 12, 2050–2080 (2017).

49. Schindelin, J. et al. Fiji: an open-source platform for biological-image analysis. Nat Meth 9, 676–682 (2012).

