## Supplementary Table, Movie Caption and Figures for "Biomolecular Contrast Agents for Optical Coherence Tomography"

### Supplementary Table, Movie and Figures

| | Halo | Ana | Mega |
|---|---|---|---|
| Molar GV concentration per $OD_{500,PS}$ (pM / $OD_{500,PS}$) | 47.3 | 114 | 2030 |
| Protein concentration per $OD_{500,PS}$ ([μg/mL] / $OD_{500,PS}$) | 13.4 | 36.6 | 145.5 |
| Estimated molecular weight (MDa) | 282 | 320 | 71.7 |
| Volumetric fraction of GV-contained gas in solution (v/v / $OD_{500,PS}$) | 0.000178 | 0.000417 | 0.000794 |

**Supplementary Table 1. Summary of the protein and gas contents of three types of gas vesicles (GVs).** GVs are usually quantified experimentally by $OD_{500,PS}$ and subsequently converted to molar concentration and GV-contained gas volume. To make the conversion, protein and gas content of GVs per $OD_{500,PS}$ were determined by a combination of transmission electron microscopy (TEM), geometric calculation and protein assays[1, 2].

**Supplementary Movie 1 | Representative OCT image series of unclustered and clustered Ana GVs.** A series of 200 images were collected over 2 seconds at 100 frames per second (fps) on unclustered (left) and clustered (right) Ana GVs (0.91 nM, $OD_{500,PS}$ = 8.0). The movie is displayed at 10 fps, 10 times slower than the real time.

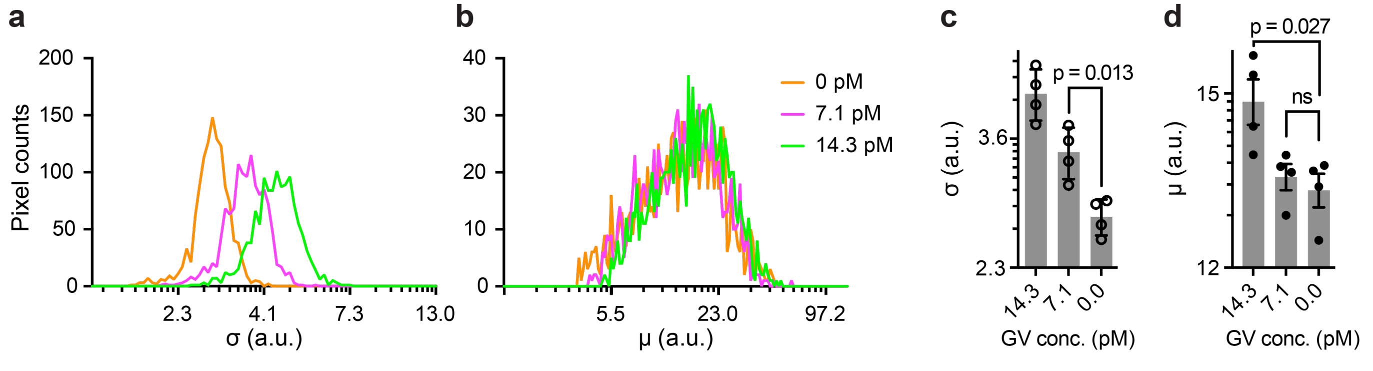


**Supplementary Figure 1 | Quantification of the image intensity of Ana GVs at low concentrations.** For the OCT images in Fig. 2d, regions of interest (ROIs) of 60 x 20 pixels were drawn for the samples containing 14.3 pM, 7.1 pM Ana GVs and the blank sample. **a-b,** Histograms of the intensity of individual pixel were plotted for the μ and σ images. **c-d**, Mean μ and σ of all the pixels in the ROIs for $N = 4$ repeats of the experiment. Unpaired *t*-test was performed and *p* values were shown.


bioRxiv preprint. March 31, 2019. All rights reserved.

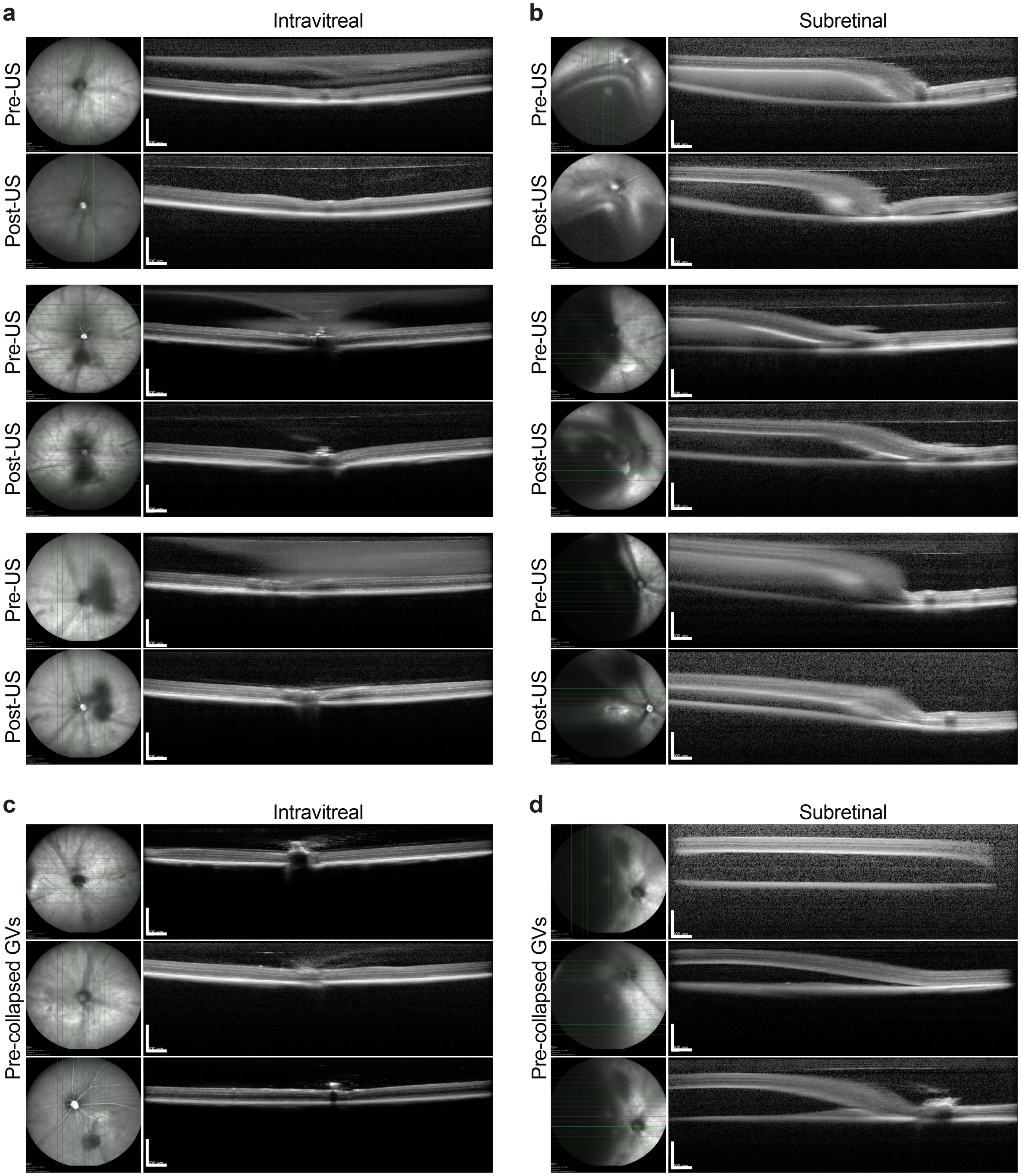


**Supplementary Figure 2 | Additional *in vivo* images of gas vesicles at mouse retina.** Fundus images and B-scan OCT images of mouse eyes intravitreally (**a, c**) and subretinally (**b, d**) injected with intact GVs before and after ultrasound treatment (**a, b**) or with GVs that were collapsed by ultrasound prior to injection (**c, d**). Scale bars represent 0.5 mm.


bioRxiv preprint. March 31, 2019. All rights reserved.

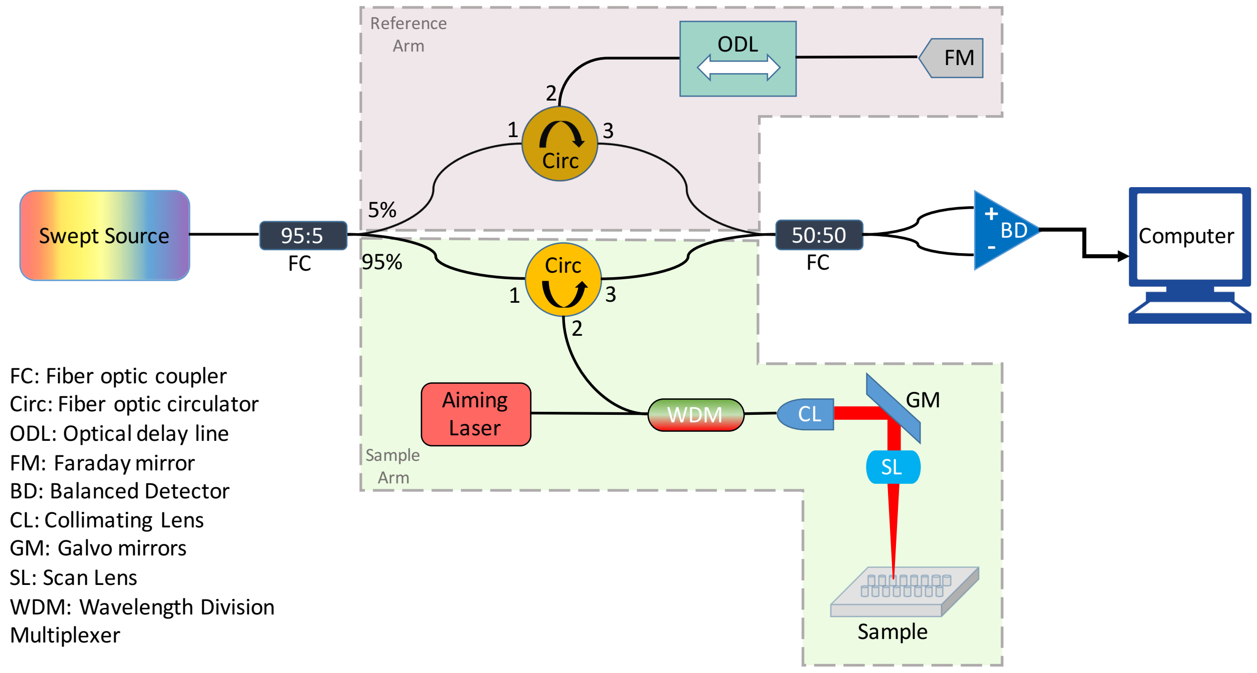


**Supplementary Figure 3 | The design and components of the custom-built OCT scanning system.** The output beam of the swept laser was split by 95:5 coupler into the sample arm and reference arm and then sent through the respective optical elements in each arm. Both the sample and reference arms contained a fiber optic circulator to route the reflected infrared beam towards the 50:50 coupler, which allowed the system to be symmetrical and maximize collection efficiency of the reflected beam from the sample target. In the sample arm, the infrared beam from port 2 of the circulator was combined with a 635-nm aiming laser beam via a wavelength division multiplexer (WDM), and then passed through a 2-axis galvo scanner and a scan lens (SL) that focused the beam onto the sample target. The red aiming laser in the system aided the visualization of the scanning area and facilitates the positioning of the sample target. The infrared beam reflected from the sample target traversed back through the same optical path, entering port 2 and exiting from port 3 of the sample arm circulator towards the 50:50 coupler. Similarly, the reference beam passed through a motorized optical delay line (ODL), was reflected by a fiber optic Faraday mirror, and subsequently mixed with the sample beam in the 50:50 coupler. The fiber optic Faraday mirror minimized changes to the polarization state of light induced by external perturbations, while the ODL was necessary to adjust and match the optical path length of the reference arm to that of the sample arm. The resolution of the OCT setup can be calculated: the 2-axis galvo setup contained a triplet fiber optic collimator (TC12APC-1310, Thorlabs) with an effective focal length of 12 mm and a scan lens (LSM03, Thorlabs) with an effective focal length of 36 mm. The fiber optic collimator output a beam with a waist diameter of 2.26 mm, and the scan lens was capable of generating a spot size of approximately 21.5 µm at the focal plane and central wavelength, assuming a 4 mm diameter input beam. Therefore, this combination of lenses generated a spot size of ~ 38 µm. In terms of axial performance, the swept laser had an approximately 96 nm sweep range, but only a range of 84 nm was sampled during data acquisition. Therefore, the theoretical axial resolution of the OCT system was about 9.3 µm in air and 7 µm in water.

### Supplementary References


1. Lu, G.J. et al. Acoustically modulated magnetic resonance imaging of gas-filled protein nanostructures. *Nature Materials* ***17***, 456–463 (2018).
2. Lakshmanan, A. et al. Preparation of biogenic gas vesicle nanostructures for use as contrast agents for ultrasound and MRI. *Nature Protocols* **12**, 2050-2080 (2017).




bioRxiv preprint. March 31, 2019. All rights reserved.
